# Transthyretin-stabilizing mutation T119M is not associated with protection against vascular disease or death in the UK Biobank

**DOI:** 10.1101/771626

**Authors:** Margaret M. Parker, Simina Ticau, James Butler, David Erbe, Madeline Merkel, Emre Aldinc, Gregory Hinkle, Paul Nioi

## Abstract

**Background:** Destabilized transthyretin (TTR) can result in the progressive, fatal disease transthyretin-mediated (ATTR) amyloidosis. A stabilizing *TTR* mutation, T119M, is the basis for a therapeutic strategy to reduce destabilized TTR. Recently, T119M was associated with extended lifespan and lower risk of cerebrovascular disease in a Danish cohort. We aimed to determine whether this finding could be replicated in the UK Biobank.

**Methods:** *TTR* T119M carriers were identified in the UK Biobank, a large prospective cohort of ∼500,000 individuals. Association between T119M genotype and inpatient diagnosis of vascular disease, cardiovascular disease, cerebrovascular disease, and mortality was analyzed.

**Results:** Frequency of T119M within the white UK Biobank population (*n*=337,148) was 0.4%. Logistic regression comparing T119M carriers to non-carriers found no association between T119M and vascular disease (odds ratio [OR]=1.08; *p=*.27), cardiovascular disease (OR=1.08; *p=*.31), cerebrovascular disease (OR=1.1; *p=*.42), or death (OR=1.2; *p=*.06). Cox proportional hazards regression showed similar results (hazard ratio>1, *p*>.05). Age at death and vascular disease diagnosis were similar between T119M carriers and non-carriers (*p=*.12 and *p=*.38, respectively).

**Conclusions:** There was no association between the *TTR* T119M genotype and risk of vascular disease or death in a large prospective cohort study, indicating that TTR tetramer stabilization through T119M is not protective in this setting.

## Introduction

Transthyretin (TTR), also known as prealbumin, is a homo-tetrameric protein that is primarily synthesized and secreted by the liver but also, to a lesser extent, by the retina and choroid plexus. TTR functions as a major transporter of the vitamin A–retinol binding protein complex, but it also serves a minor role in the transport of thyroxine [1–3].

In cases where the *TTR* gene is mutated or a spontaneous destabilization of the protein occurs, TTR tetramers can dissociate into monomers. These monomers can misfold into amyloid fibrils and form insoluble deposits in multiple tissues including the heart, peripheral nerves, and gastrointestinal tract [4, 5]. Additionally, although less well described, TTR can deposit in cerebral blood vessels (i.e., cerebral amyloid angiopathy) and associated hemorrhagic and ischemic lesions have been noted in a small number of patients [6–9]. The deposition of TTR amyloid in multiple organs and tissues can result in transthyretin-mediated (ATTR) amyloidosis, a progressively debilitating and life-threatening disease [5,10,11]. Two types of ATTR amyloidosis exist: (1) wild-type ATTR amyloidosis, which occurs when no *TTR* mutation is present and predominantly manifests as cardiomyopathy (although other systems can be involved); and (2) hereditary transthyretin-mediated (hATTR) amyloidosis, where over 120 distinct pathogenic mutations have been identified that serve to destabilize tetrameric TTR, leading to dissociation and subsequent fibril deposition [12, 13]. hATTR amyloidosis results in multisystem disease that can include sensory and motor, autonomic, and cardiac symptoms [14].

Two mutations encoding a thermodynamically and kinetically stabilized TTR protein have also been described, *TTR* Thr119Met (T119M; rs28933981) and *TTR* Arg104His (R104H; rs121918095) [15–18]. Both variants were discovered in heterozygotes carrying a pathogenic *TTR* variant in *trans* with a stabilizing variant, resulting either in a milder pathology or no detectable disease compared with carriers of pathogenic *TTR* variants alone [19–21]. When these stabilizing variants are present in combination with pathogenic TTR, the resulting hetero-tetramers demonstrate increased stability *in vitro* [16,18,22]. Specifically, T119M is thought to slow the dissociation of the TTR tetramer by increasing the hydrophobic surface area at the dimer/dimer interface [23]. These observations inspired the discovery of small-molecule TTR tetramer stabilizing therapeutics [24].

While a few case studies have suggested that T119M and R104H protect against severe manifestations of hATTR amyloidosis, it is not clear whether they have any effect in non-carriers of pathogenic *TTR* mutations. It has been reported that T119M carriers have higher circulating levels of TTR compared with non-carriers, but no phenotypic associations have been noted for this variant in the genome-wide association catalog despite it being carried by ∼1 in 200 people [25, 26]. A previous publication in a Danish population cohort suggested that carriers of T119M may be protected against cerebrovascular disease and have longer lifespan [25]. However, these findings have not been replicated to date.

In the current study, we investigated the potential effect of *TTR* T119M on vascular disease and mortality in the UK Biobank cohort to assess whether previously documented findings could be replicated.

## Methods

### Study population

The UK Biobank is a population-based prospective cohort study that recruited ∼500,000 participants aged 40–69 years in England, Wales, and Scotland between 2006 and 2010. Participants had to live within 25 miles of one of the 22 assessment centers and participate in a baseline assessment involving the collection of extensive data from questionnaires, health records, physical measurements, imaging, and biologic samples [27]. This analysis included all UK Biobank participants classified as unrelated, white British based on a combination of self-reported ancestry and genetic principal components (*n*=337,148) [28]. Data used in this analysis were accessed through UK Biobank application 26,041 and are publicly available upon request to the UK Biobank. The UK Biobank holds ethical approval and all participants provided written informed consent.

### T119M genotyping

Details of the genotyping and quality control process of genetic data in the UK Biobank have been described elsewhere [29]. Briefly, UK Biobank participants were genotyped on either the UK BiLEVE or UK Biobank Axiom array, which share >95% common content. Genotypic data were imputed to the UK10K haplotype, 1000 Genomes Phase 3, and Haplotype Reference Consortium reference panels using SHAPEIT3 [30] for phasing and IMPUTE4 [29] for imputation. The T119M genotype (rs28922981) was imputed with high accuracy (info score=0.925). For cross-sectional analyses, T119M genotype was included in the statistical model as dosages, and for analyses requiring a hard-called genotype, the best-guessed genotype was used.

### Ascertainment of disease and death

We analyzed a composite phenotype of vascular disease, defined as a primary or secondary ICD10 diagnoses of I20–I25, I60–I69, or G45, including all child codes of these diagnoses. Additionally, the components of vascular disease (cardiovascular [CV] disease [I20–I25] and cerebrovascular disease [I60–I69 or G45]) as well as the sub-components of cerebrovascular disease (ischemic cerebrovascular disease [I64–I69 and G45]) and hemorrhagic stroke (I60–I63) were also separately analyzed. All ICD10 diagnoses were taken from field 41270 from the UK Biobank data release on November 3, 2019. ICD10 codes were collected through March 31, 2017, for participants in England, February 29, 2016, for participants in Wales, and October 31, 2016, for participants in Scotland.

Date of death (field 40,000) was obtained from linkage to the National Death Registries. All data from death certificates are obtained by the UK Biobank on a quarterly basis. Data on death were collected through January 31, 2018, for participants in England and Wales and November 30, 2016, for participants in Scotland.

### Covariates

Participant age (field 21,002), sex (field 22,001), and smoking status (field 20,116) were based on self-report at time of enrollment. Body mass index (BMI) (field 21,001) was measured as weight in kilograms divided by height in meters squared (kg/m^2^) at time of enrollment. Hypertension was defined as a primary or secondary ICD10 diagnosis (field 41,270) of essential hypertension (I10). Diabetes was defined as a primary or secondary ICD10 diagnosis (field 41,270) of diabetes (E11). Participants were verbally interviewed by a trained nurse about medication use (field 20,003). Lipid-lowering therapy was defined as reported use of simvastatin, atorvastatin, eptastatin, fluvastatin, rosuvastatin, pravastatin, atorvastatin, velastatin, Lipitor, Zocor, Lescol, Crestor, niacin, fibrates, or ezetimibe. A total of 34 serum biomarkers were measured centrally by the UK Biobank using blood drawn at the time of enrollment. This biomarker panel provided measurements for each participant of triglycerides in mmol/L (field 30,870), low-density lipoprotein (LDL) cholesterol in mmol/L (field 30,780), high-density lipoprotein (HDL) cholesterol in mmol/L (field 30,760) and C-reactive protein (CRP) in mg/L (field 30,710).

### Statistical analysis

Descriptive statistics are presented as means (standard deviation) for continuous variables and percentages for categoric variables. Continuous and categoric variables were compared between T119M carriers and non-carriers using a t-test and a Pearson chi-square test, respectively.

The association of T119M genotype to death and vascular disease (a composite of CV and cerebrovascular diagnoses) was analyzed using logistic regression and Cox proportional hazards models. Participants with vascular diagnoses prior to UK Biobank study entry, defined as ICD10 diagnosis of disease between 1996 and enrollment date, were removed from the prospective time to vascular diagnosis analyses. For mortality analyses, time on study was calculated as time from the date of enrollment to date of death (if participant died), date of lost to follow-up (if participant was lost to follow-up), or last date of data extraction from National Death Registries (January 31, 2018, for participants in England and Wales or November 30, 2016, for participants in Scotland).

For Cox analyses with ICD10 outcomes, time on study was calculated as time from the date of enrollment to date of first vascular diagnosis (if participant had a vascular diagnosis), date of lost to follow-up (if participant was lost to follow-up), date of death (if participant died), or administrative censoring date (March 31, 2017, for participants in England, October 31, 2016, for participants in Scotland, or February 29, 2016, for participants in Wales). Data availability dates were obtained from: http://biobank.ctsu.ox.ac.uk/crystal/exinfo.cgi?src=Data_providers_and_dates (accessed on: April 1, 2019). Kaplan–Meier curves were used to estimate survival to first vascular diagnosis or death by T119M genotype. Because six outcomes were tested (death, vascular disease, cerebrovascular disease, CV disease, ischemic cerebrovascular disease, and hemorrhagic stroke), a *p*-value of .008 (.05/6) was considered as statistically significant. Survival analyses were performed in R version 3.4.4 using the survival (version 2.43) and survminer packages (version 0.4.3). All analyses were controlled for known confounders including age, sex, smoking status, BMI, and genetic ancestry via 10 genetic principal components. Additionally, a sensitivity analysis was performed controlling for the available confounders presented in Hornstrup et al. [25] (age, sex, smoking status, BMI, triglycerides, LDL, HDL, hypertension, diabetes, CRP, lipid-lowering therapy use, and genetic ancestry). Analyses were performed in PLINK (version 2.0b) using the REVEAL/SciDB translational analytics platform from Paradigm4.

Age at death or diagnosis of first vascular disease was compared between T119M and non-carriers using a t-test. For participants that were diagnosed with a vascular disease and subsequently died, the number of years they survived post diagnosis was compared using a t-test.

## Results

### Baseline characteristics

**Table 1** shows the baseline characteristics of the 337,148 participants included in this study. A total of 2502 UK Biobank participants carried the T119M allele, distributed as 2499 heterozygotes and three homozygotes (minor allele frequency=0.4%). T119M carriers were less likely to be smokers than non-carriers (*p=*.02) but the two groups were similar in terms of age, sex, and other CV risk factors (**Table 1**).

**Table 1.** Characteristics of UK Biobank study population by T119M genotype.

|  | Non-carriers<br>(CC)<br>n=334,646 | Carriers<br>(CT or TT)<br>n=2502 | *p*-value |
| --- | --- | --- | --- |
| Mean (SD) age, years | 56.9 (8.0) | 56.7 (8.0) | .16 |
| Male (%) | 46.3 | 47.0 | .50 |
| Mean (SD) triglycerides (mmol/L) | 1.8 (1.0) | 1.8 (1.1) | .81 |
| Mean (SD) LDL, mmol/L | 3.6 (0.9) | 3.5 (0.9) | .14 |
| Mean (SD) HDL, mmol/L | 1.5 (0.4) | 1.5 (0.4) | .36 |
| Mean (SD) BMI, kg/m<sup>2</sup> | 27.4 (4.8) | 27.4 (4.7) | .87 |
| Mean (SD) C-reactive protein, mg/L | 2.6 (4.4) | 2.5 (4.4) | .43 |
| Hypertension (%) | 22.2 | 21.0 | .14 |
| Diabetes (%) | 5.3 | 5.7 | .40 |
| Smoking (%) | 45.2 | 42.8 | .02 |
| Lipid-lowering therapy (%)<sup>a</sup> | 18.6 | 17.9 | .43 |

<sup>a</sup>Lipid-lowering therapy was defined as self-reported use of atorvastatin, Crestor, eptastatin, ezetimibe, fibrate, fluvastatin, Lipitor, Lescol, niacin, pravastatin, simvastatin, rosuvastatin, velastatin, or Zocor.
BMI: body mass index; HDL: high-density lipoprotein; LDL: low-density lipoprotein; SD: standard deviation.

### Association of TTR T119M genotype with vascular disease and death

The association of T119M genotype with death and vascular disease was investigated using logistic regression controlling for age, sex, smoking status, BMI, and genetic ancestry. There was a total of 13,680 deaths and 36,024 vascular disease diagnoses captured in the UK Biobank. No significant association was identified with any of the outcomes examined, namely, death, all vascular disease, CV disease, cerebrovascular disease, ischemic vascular disease, or hemorrhagic stroke (**Figure 1**), or with any of the constituent diseases that these compound outcomes are composed of (**Supplemental Table S1**). Similarly, no significant association of T119M genotype was identified with any outcome examined after additionally controlling for hypertension, diabetes, LDL, HDL, triglycerides, and CRP (**Supplemental Table S2**).

**Figure 1.**
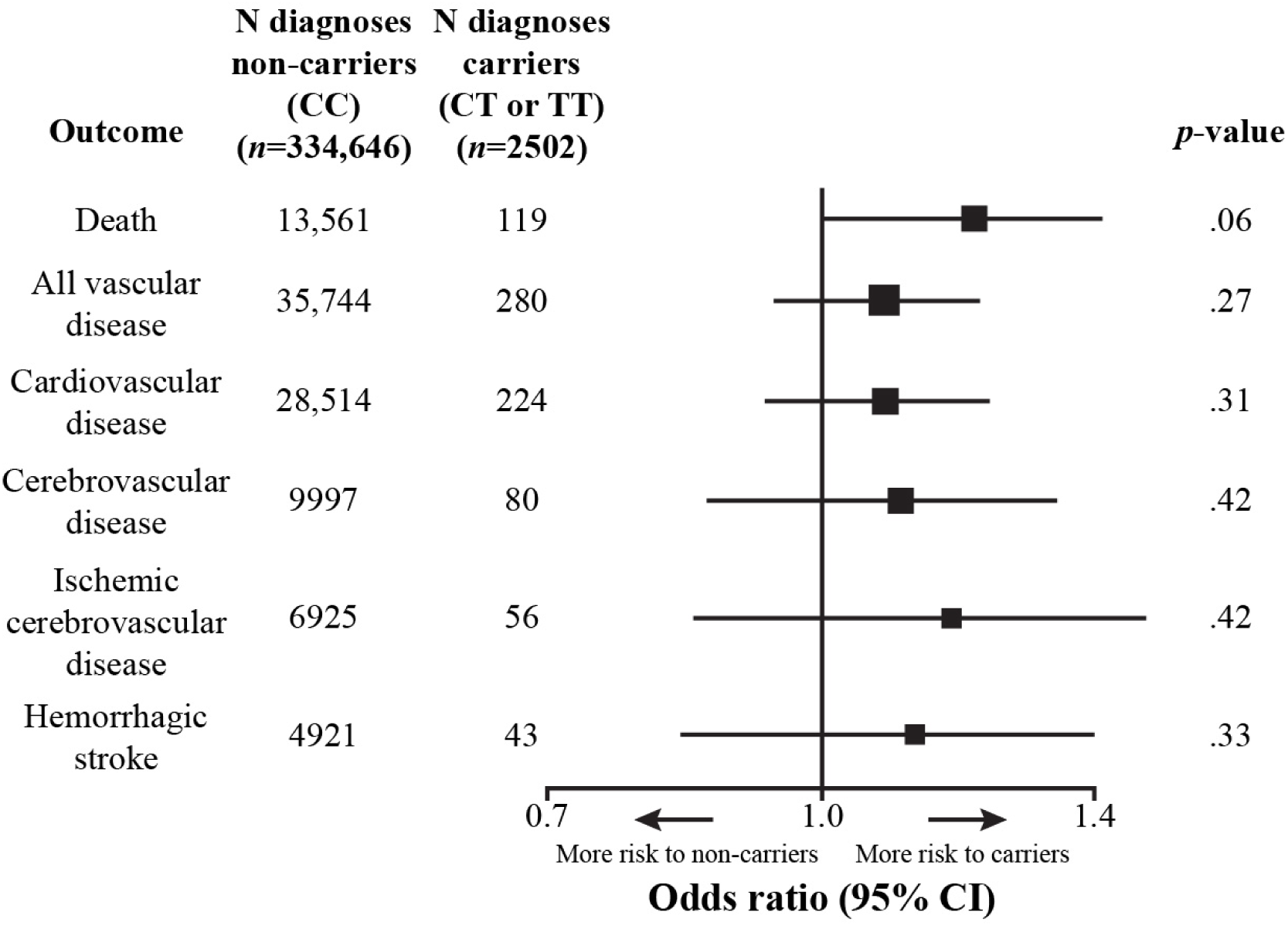
Association of vascular disease, cardiovascular disease, cerebrovascular disease, ischemic cerebrovascular disease, or hemorrhagic stroke as a function of T119M genotype using logistic regression. Odds ratios are represented by squares proportional to sample size. Genotypes = CC: T119M non-carrier; CT: T119M heterozygous carrier; TT: T119M homozygous carrier. CI: confidence interval.

### TTR T119M genotype and risk of vascular disease and death in time to diagnosis analysis

In addition to the cross-sectional analysis, a Cox proportional hazards analysis was used to investigate whether an association exists between the T119M genotype and risk of death or vascular disease. On average, UK Biobank participants were followed for 8.9 years (range: 0–11.9 years) and were 65.7 years old at censoring (range: 40.6–81.1 years) (**Supplemental Figures S1 and S2**). Loss to follow-up over the study period was minimal (0.2%) and did not differ by T119M genotype (*p=*.5). Prevalent diagnoses occurring before a patient’s date of enrollment were removed, resulting in 19,804 vascular disease diagnoses and 13,680 deaths over a total of 2,959,979 person-years of follow-up. No deaths or vascular disease diagnoses occurred in the three T119M homozygotes. Cox proportional hazards regression controlling for age, sex, BMI, smoking, and genetic ancestry revealed no difference between T119M carriers and non-carriers in their risk of death or vascular disease, including CV, cerebrovascular, hemorrhagic stroke, and ischemic cerebrovascular disease (**Figure 2**). This also remained true after controlling for hypertension, diabetes, LDL, HDL, triglycerides, and CRP (**Supplemental Table S3**). Vascular disease diagnoses occurred in 6.2% of T119M carriers and 5.9% of non-carriers; survival time to first vascular disease diagnosis was not lengthened in T119M carriers (HR=1.06; 95% CI = 0.8–1.2; *p=*.45) (**Figure 3**).

**Figure 2.**
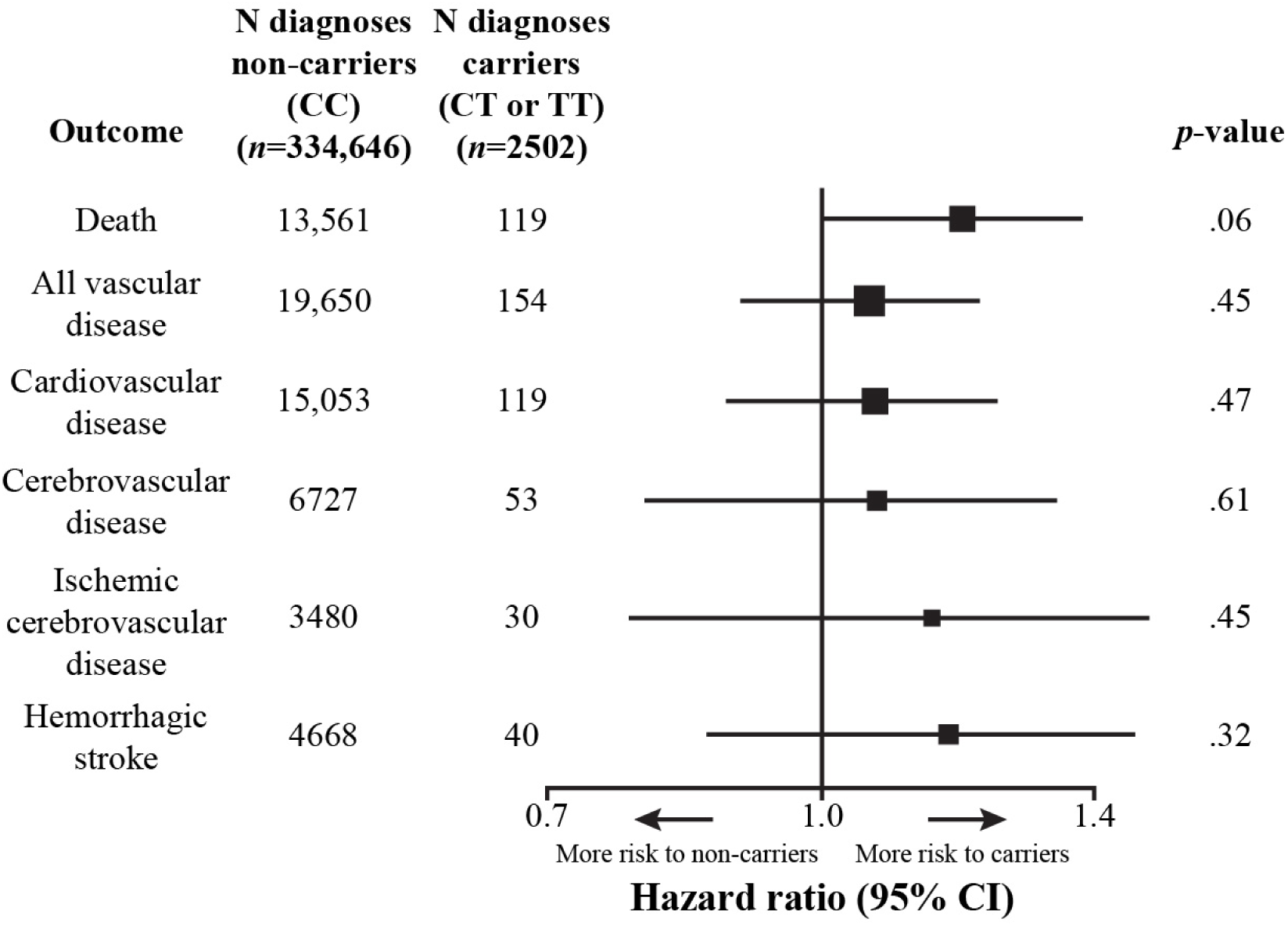
Risk of vascular disease, cardiovascular disease, cerebrovascular disease, ischemic cerebrovascular disease, or hemorrhagic stroke as a function of T119M genotype using Cox proportional hazard analysis. Hazard ratios are represented by squares proportional to sample size. Genotypes = CC: T119M non-carrier; CT: T119M heterozygous carrier; TT: T119M homozygous carrier. CI: confidence interval.

**Figure 3.**
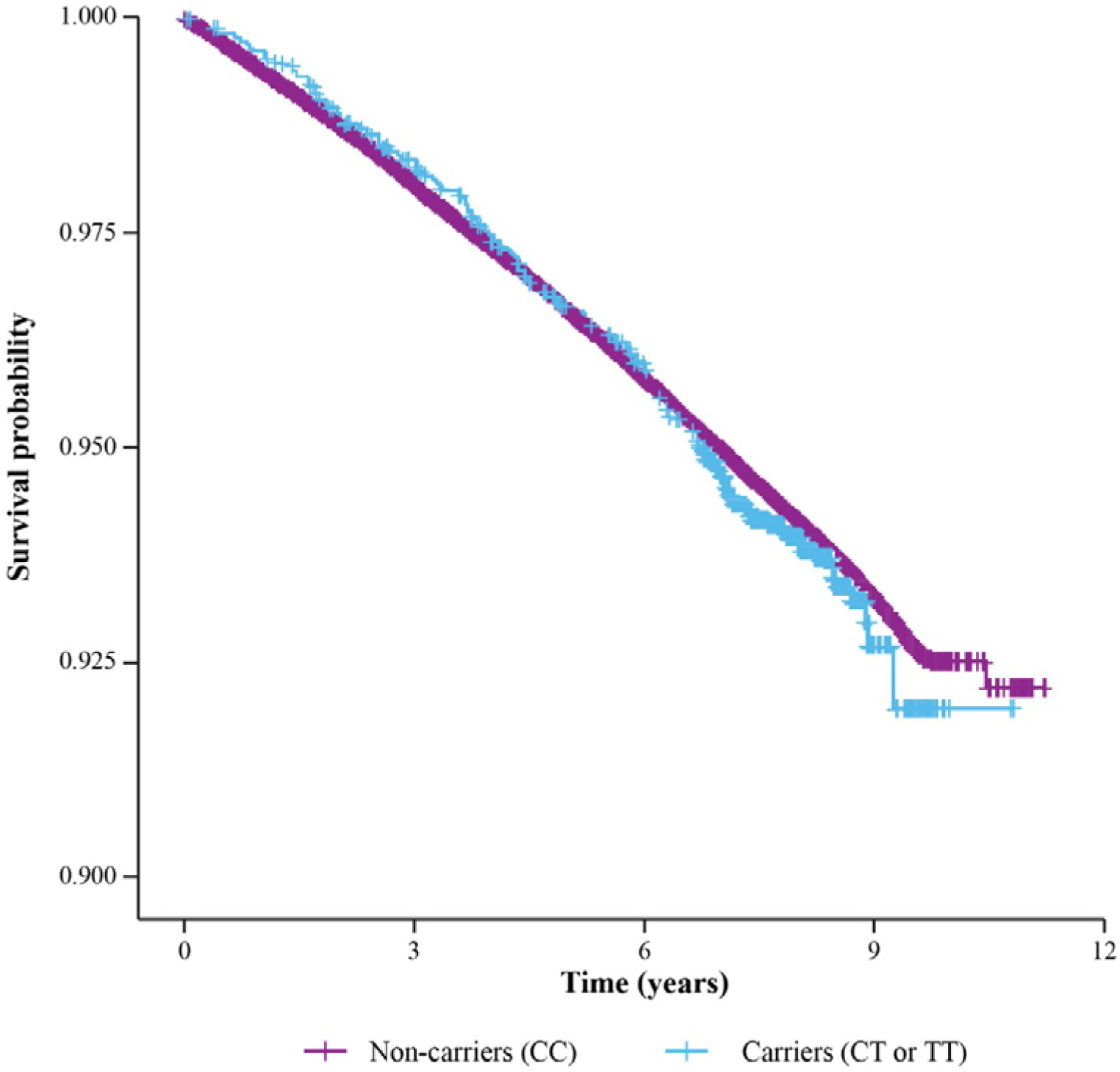
Time to first vascular disease diagnosis by T119M genotype. In Cox proportional hazard analysis controlling for age, sex, smoking status, BMI and genetic ancestry, there was no association between T119M genotype and time to first vascular disease diagnosis (HR=1.06, *p=*.45). Genotypes = CC: T119M non-carrier; CT: T119M heterozygous carrier; TT: T119M homozygous carrier. BMI: body mass index.

### T119M genotype and age at death and survival following diagnosis of vascular disease

The mean age at death from all causes was similar between T119M carriers and non-carriers (*p=*.12) and no difference was seen in age at first diagnosis of vascular disease (*p=*.38), including cardiovascular disease (*p=*.72) and cerebrovascular disease (*p=*.41) (**Figure 4**). For participants who were diagnosed with any vascular disease and subsequently died (*n*=4180), there was no significant difference between T119M carriers and non-carriers in their survival time following vascular disease diagnosis (*p=*.82), including CV disease (*p=*.84) and cerebrovascular disease (*p=*.41) (**Table 2**).

**Figure 4.**
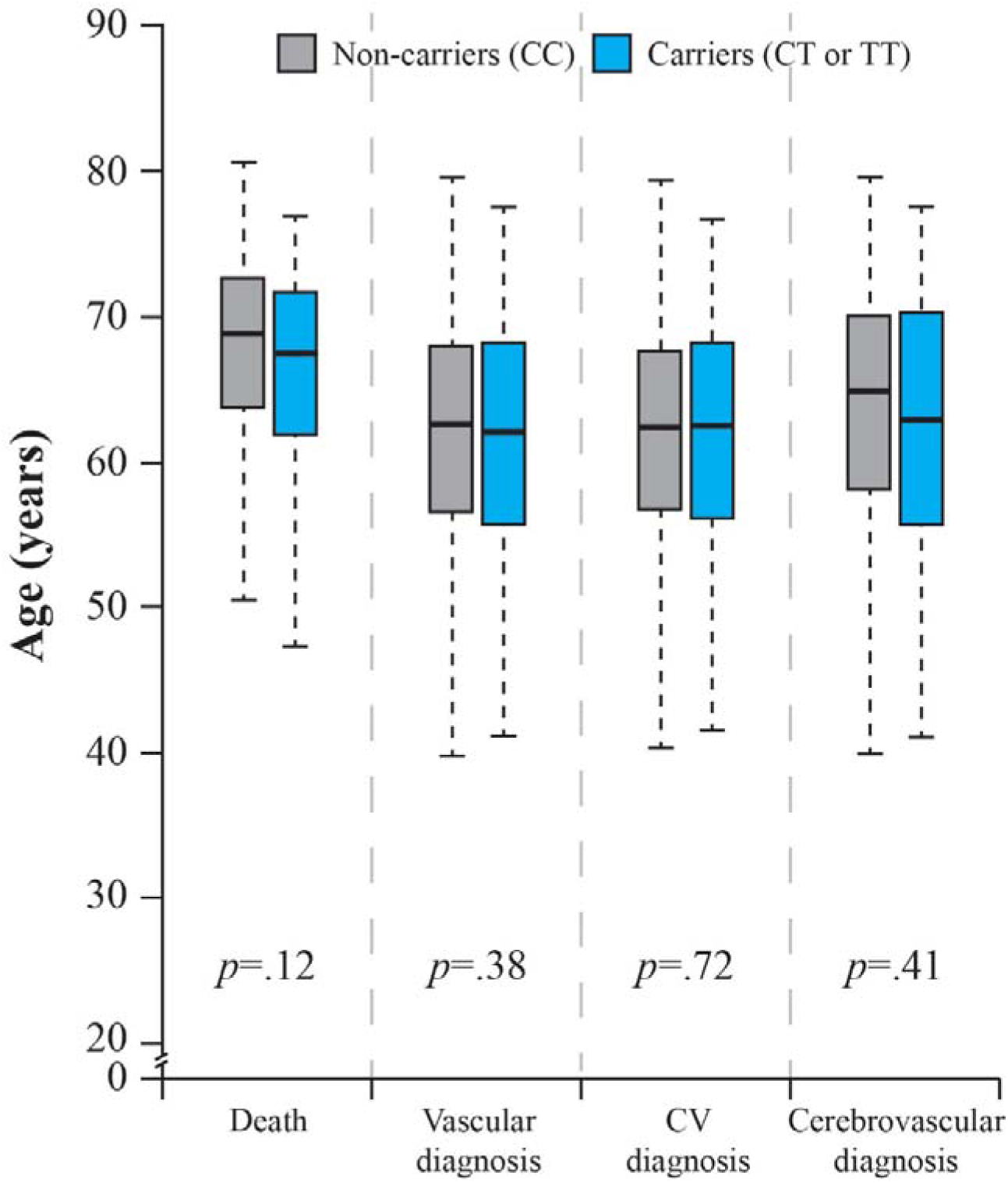
Age at death or diagnosis by T119M genotype. There is no difference between T119M carriers and non-carriers in age at death (t-test *p=*.12), age at first diagnosis of vascular disease (t-test *p=*.38), age at first diagnosis of CV disease (t-test *p=*.72), or age at first diagnosis of cerebrovascular disease (t-test *p=*.41). Genotypes = CC: T119M non-carrier; CT: T119M heterozygous carrier; TT: T119M homozygous carrier. CV: cardiovascular.

**Table 2.**
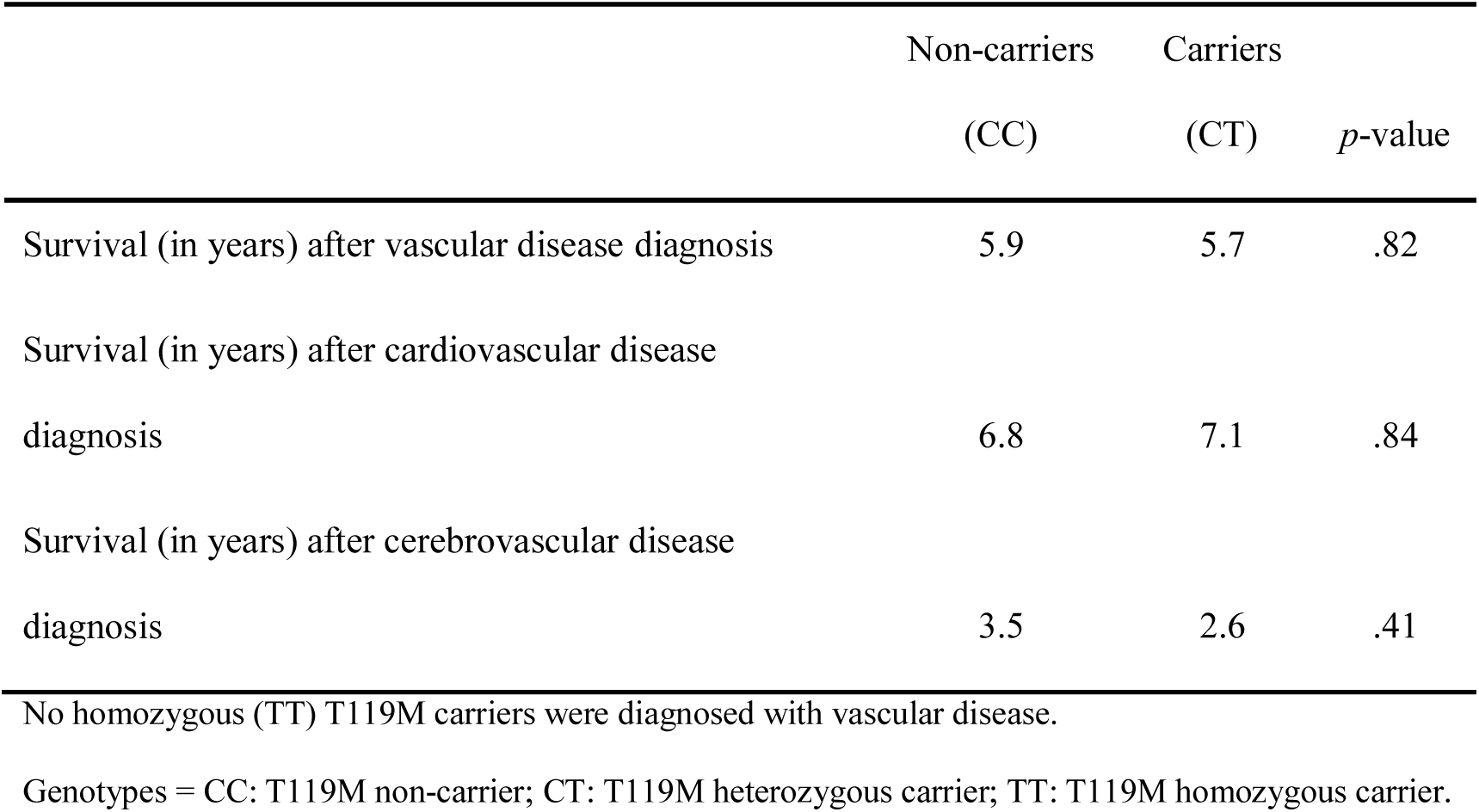
Mean survival (in years) after diagnosis of vascular, cardiovascular, or cerebrovascular disease. There was no difference in time to death for those who were diagnosed with any vascular disease, cardiovascular disease, or cerebrovascular disease and subsequently died.

## Discussion

In a large prospective cohort study of 337,148 participants of the UK Biobank, carriers of the *TTR* T119M mutation were not found to be protected against vascular disease, cerebrovascular disease, CV disease, or death (*p*>.05, OR/HR >1 for all analyses). Furthermore, no difference was seen between T119M carriers and non-carriers in their time to death following a diagnosis of vascular disease (including CV and cerebrovascular disease). Together, these findings suggest stabilization of the TTR tetramer via the T119M mutation is not beneficial in a general population setting in terms of mortality and vascular disease.

These findings are notable given a previous report from a Danish cohort showing T119M carriers were protected against vascular disease and have longer lifespans than non-carriers [25]. In that study, the authors compared 321 T119M heterozygotes to ∼68,000 non-carriers in a cohort containing 10,636 incident vascular diagnoses spanning an average follow-up of 32 years. The current study offered a larger sample size (2502 T119M carriers and 334,646 non-carriers) and, therefore, greater statistical power to detect associations between T119M genotype and vascular disease or death. This current study has 98% power to detect an association with vascular disease with OR of 0.83 compared with 52% power in the study by Hornstrup et al. [25, 31]. Overall, the UK Biobank includes 7.8 times more T119M carriers (2502 vs. 321) and 3.4 times more vascular diagnoses. Moreover, the study by Hornstrup et al. [25] had a small number of diagnoses within the T119M carriers (*n*=37 vascular diagnoses, 29 cardiovascular diagnoses, and 11 cerebrovascular diagnoses). Additionally, there are other methodologic differences that may explain the contrasting results. In the present study, related subjects were removed from the analyses, and genetic principal components were used to control for population stratification in the regression models, both of which are factors which, if not accounted for, can lead to false positive results. The mean follow-up time of individuals was different between the studies (mean of 8.9 years in the current study versus 32 years in the previous report). However, given there are 7.8 times more carriers and 3.4 times more diagnoses in this analysis, it is not clear whether length of follow-up would be expected to provide an explanation for the observed differences. Finally, geographic differences, including diet, lifestyle, and climate, may also have an effect as these two studies were performed in different populations.

These results raise questions concerning what, if any, physiologic effects the *TTR* T119M mutation confers to the general population. However, the biochemical effects of this variant are clear; threonine 119 sits within the deepest of the three halogen binding pockets of the T4 binding site created by the dimer–dimer interface in TTR. The substitution of methionine at this position introduces additional hydrophobicity in the pocket, inducing additional contacts between the different dimers at this interface, resulting in enhanced kinetic stability [15–18,21]. In the context of the carriers of hATTR amyloidosis-associated *TTR* gene mutations, compound heterozygotes of pathogenic *TTR* variants and the T119M variant displayed no or mild clinical symptoms of hATTR amyloidosis in a small case study, indicating a potential protective effect of the T119M mutation in this group [20]. Additionally, as a therapeutic strategy in patients with ATTR amyloidosis, TTR stabilizers mimicking the T119M mutation have shown clinical benefits [32, 33]. However, a growing body of literature indicates that patients treated with TTR stabilizer therapies continue to experience disease progression [34–40] indicating that TTR stabilization (at least with current molecules) may not be enough to stabilize or halt disease.

In summary, no association was found between T119M genotype and risk of vascular disease or death in a large-scale prospective cohort study of 337,148 participants. This suggests that stabilization of TTR through the T119M mutation does not confer general protection against vascular disease or death. Further research is needed to understand the role of TTR stabilization in ATTR amyloidosis pathogenesis as there is growing interest in treating patients with ATTR amyloidosis, including earlier intervention in identified carriers of pathogenic *TTR* mutations.

## Acknowledgments

We thank the staff and participants of the UK Biobank study. This research has been conducted under UK Biobank application 26041.

## Supplementary material

**Supplemental Table S1.** Cross-sectional association analysis of subjects with versus without the of T119M genotype and ICD10 codes that define vascular disease. Association was tested using logistic regression controlling for age, sex, smoking status, body mass index, and genetic ancestry via principal components. Codes with at least 100 diagnoses were tested for association.

| Phenotype | Odds ratio | Standard error | *p*-value |
| --- | --- | --- | --- |
| I20 | 1.02 | 0.10 | .86 |
| I21 | 1.29 | 0.13 | .05 |
| I22 | 0.98 | 0.52 | .96 |
| I24 | 0.69 | 0.37 | .32 |
| I25 | 1.06 | 0.08 | .46 |
| I60 | 0.53 | 0.62 | .30 |
| I61 | 0.94 | 0.47 | .89 |
| I62 | 1.37 | 0.55 | .56 |
| I63 | 1.27 | 0.20 | .22 |
| I64 | 0.75 | 0.47 | .54 |
| I65 | 1.28 | 0.36 | .49 |
| I66 | 1.87 | 1.03 | .54 |
| I67 | 1.13 | 0.23 | .62 |
| I69 | 1.87 | 0.27 | .02 |
| G45 | 1.22 | 0.23 | .38 |

**Supplemental Table S2.**
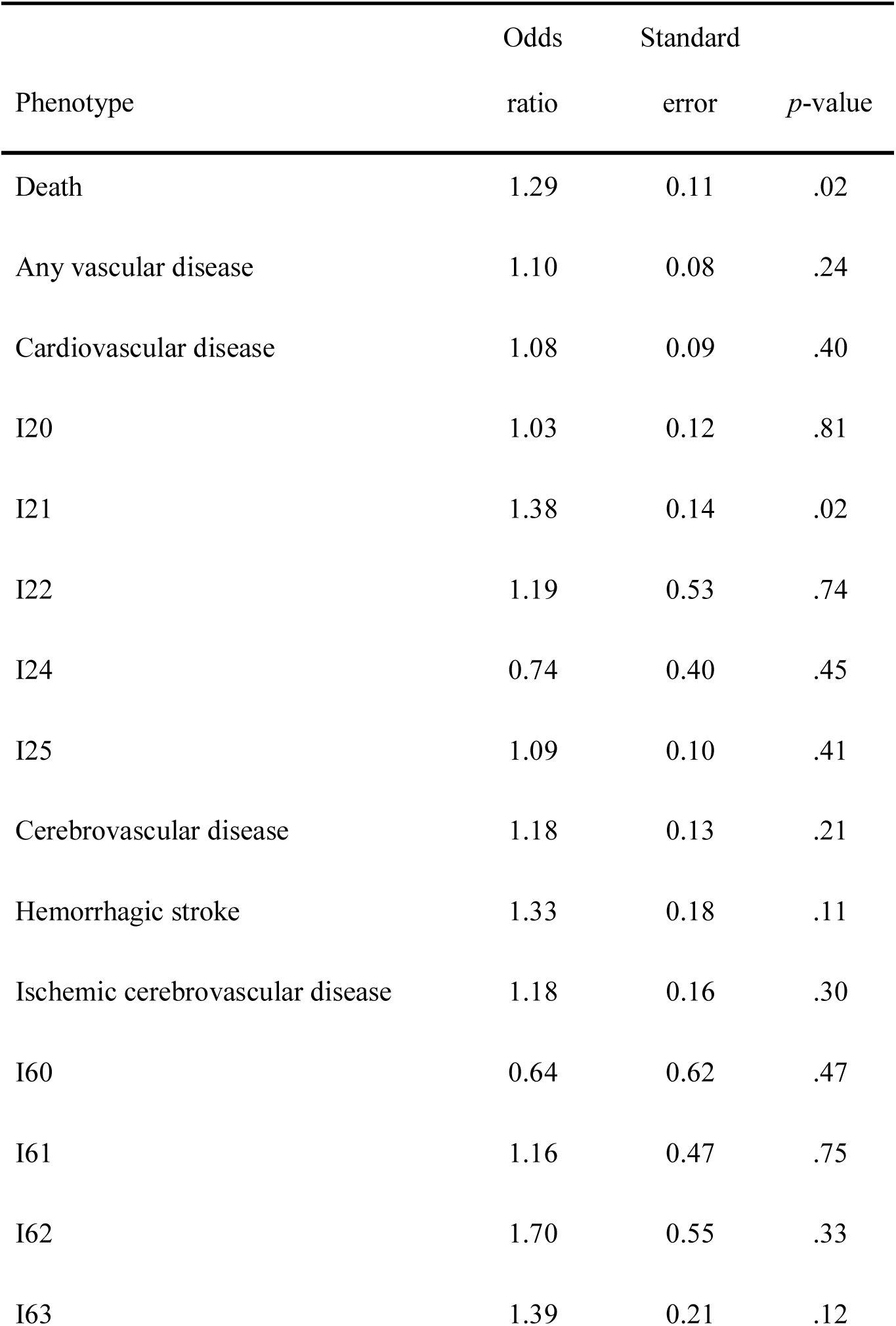
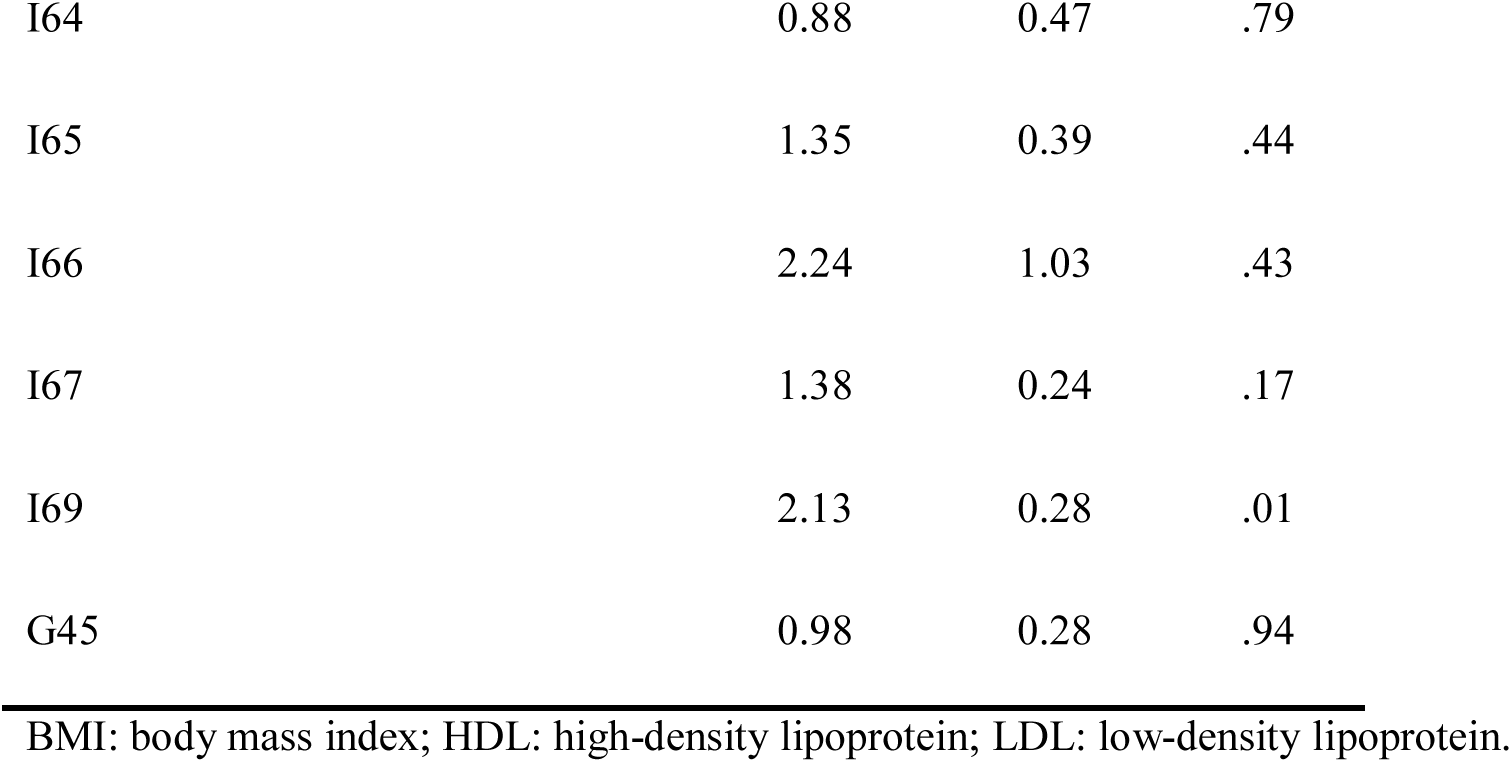
Cross-sectional association analysis of subjects with versus without the T119M genotype and all disease outcomes controlling for age, sex, BMI, smoking status, hypertension, diabetes, LDL cholesterol, HDL cholesterol, triglycerides, C-reactive protein, use of lipid-lowering drugs, and genetic ancestry via principal components. Association was tested using logistic regression. Codes with at least 100 diagnoses were tested for association.

| Phenotype | Odds<br>ratio | Standard<br>error | *p*-value |
| --- | --- | --- | --- |
| Death | 1.29 | 0.11 | .02 |
| Any vascular disease | 1.10 | 0.08 | .24 |
| Cardiovascular disease | 1.08 | 0.09 | .40 |
| I20 | 1.03 | 0.12 | .81 |
| I21 | 1.38 | 0.14 | .02 |
| I22 | 1.19 | 0.53 | .74 |
| I24 | 0.74 | 0.40 | .45 |
| I25 | 1.09 | 0.10 | .41 |
| Cerebrovascular disease | 1.18 | 0.13 | .21 |
| Hemorrhagic stroke | 1.33 | 0.18 | .11 |
| Ischemic cerebrovascular disease | 1.18 | 0.16 | .30 |
| I60 | 0.64 | 0.62 | .47 |
| I61 | 1.16 | 0.47 | .75 |
| I62 | 1.70 | 0.55 | .33 |
| I63 | 1.39 | 0.21 | .12 |
| I64 | 0.88 | 0.47 | .79 |
| I65 | 1.35 | 0.39 | .44 |
| I66 | 2.24 | 1.03 | .43 |
| I67 | 1.38 | 0.24 | .17 |
| I69 | 2.13 | 0.28 | .01 |
| G45 | 0.98 | 0.28 | .94 |

BMI: body mass index; HDL: high-density lipoprotein; LDL: low-density lipoprotein.

**Supplemental Table S3.** Risk of death, vascular disease, cardiovascular disease, cerebrovascular disease, hemorrhagic stroke, and ischemic cerebrovascular disease by T119M genotype. Association was tested using Cox proportional hazard regression controlling for age, sex, BMI, smoking status, hypertension, diabetes, LDL cholesterol, HDL cholesterol, triglycerides, C-reactive protein, use of lipid-lowering drugs, and genetic ancestry via principal components

| Phenotype | Odds<br>ratio | Standard<br>error | *p*-value |
| --- | --- | --- | --- |
| Death | 1.26 | 0.10 | .02 |
| Any vascular disease | 1.01 | 0.09 | .93 |
| Cardiovascular disease | 1.00 | 0.10 | .99 |
| Cerebrovascular disease | 1.08 | 0.15 | .59 |
| Hemorrhagic stroke | 1.23 | 0.20 | .29 |
| Ischemic cerebrovascular disease | 1.19 | 0.17 | .32 |

BMI: body mass index; HDL: high-density lipoprotein; LDL: low-density lipoprotein.

**Supplemental Figure S1.**
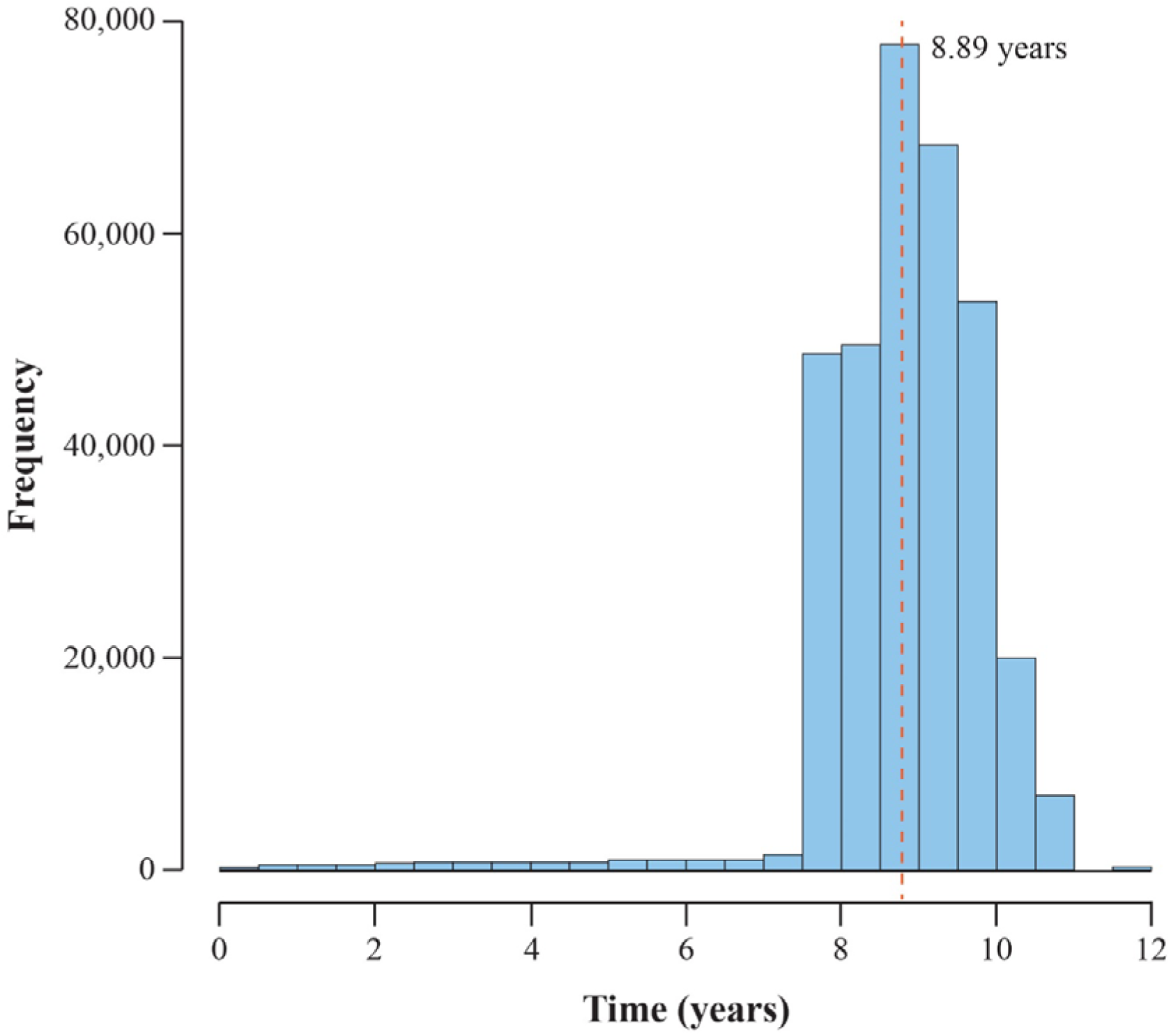
Histogram of follow-up time in the UK Biobank study. Participants were followed for an average of 8.89 years (interquartile range: 8.27–9.47 years).

**Supplemental Figure S2.**
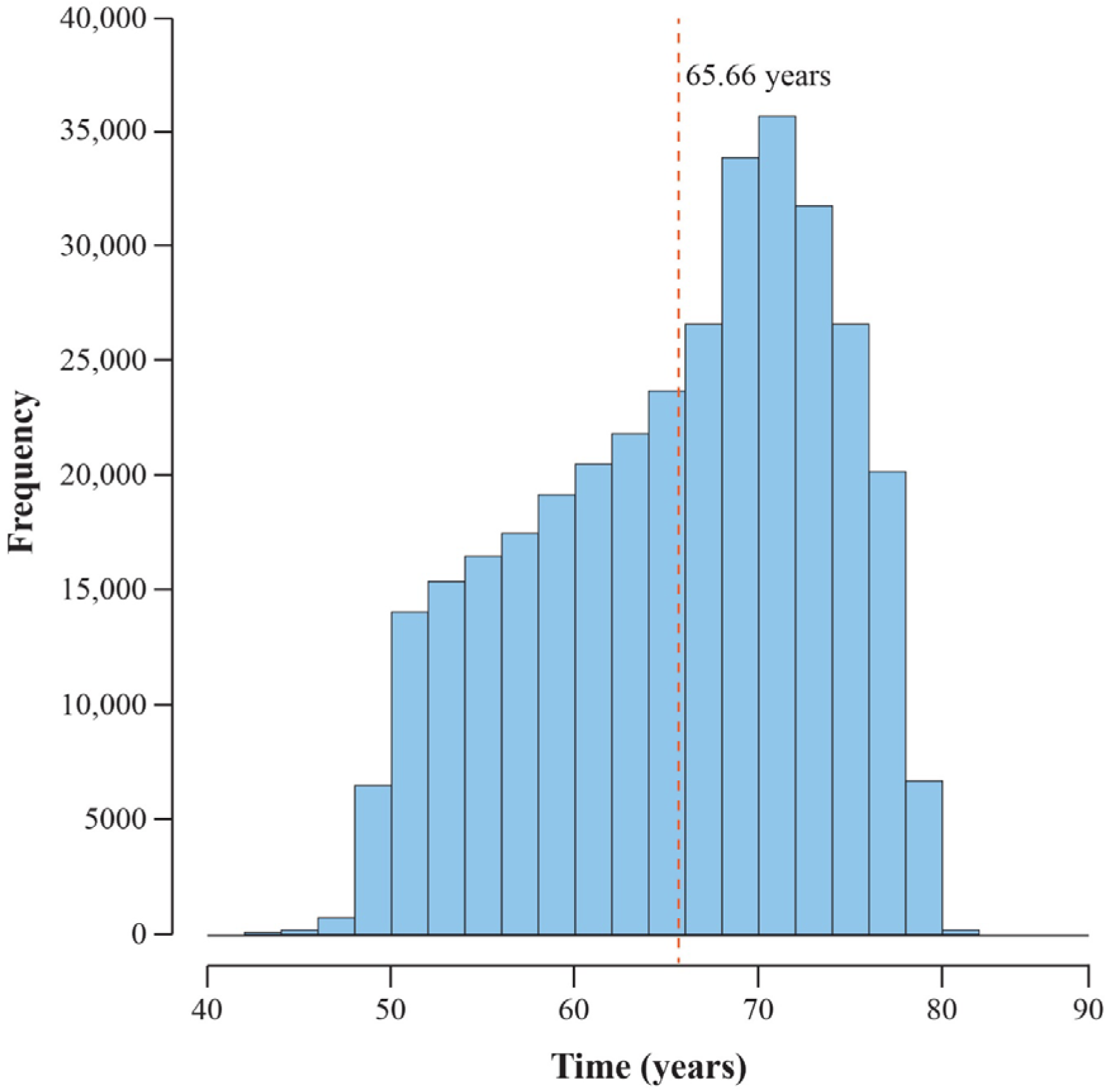
Histogram of age at last known UK Biobank observation. Participants were on average 65.66 years old at censoring (interquartile range: 59.5–72.1 years). Censoring occurred at date of death (if died), date of loss to follow-up (if lost to follow-up), or March 31, 2017 for participants in England, February 29, 2016 for participants in Wales or October 31, 2016 for participants in Scotland.

## References

[1] Cavallaro T, Martone RL, Dwork AJ, et al. The retinal pigment epithelium is the unique site of transthyretin synthesis in the rat eye. Invest Ophthalmol Vis Sci. 1990;31:497–501.

[2] Holmgren G, Steen L, Ekstedt J, et al. Biochemical effect of liver transplantation in two Swedish patients with familial amyloidotic polyneuropathy (FAP-met^30^). Clin Genet. 1991;40:242–246.

[3] Soprano DR, Herbert J, Soprano KJ, et al. Demonstration of transthyretin mRNA in the brain and other extrahepatic tissues in the rat. J Biol Chem. 1985;260:11793–11798.

[4] Damy T, Judge DP, Kristen AV, et al. Cardiac findings and events observed in an open-label clinical trial of tafamidis in patients with non-Val30Met and non-Val122Ile hereditary transthyretin amyloidosis. J Cardiovasc Trans Res. 2015;8:117–127.

[5] Hanna M. Novel drugs targeting transthyretin amyloidosis. Curr Heart Fail Rep. 2014;11:50– 57.

[6] Pitkänen P, Westermark P, Cornwell GG, 3rd. Senile systemic amyloidosis. Am J Pathol. 1984;117:391–399.

[7] Westermark P, Bergström J, Solomon A, et al. Transthyretin-derived senile systemic amyloidosis: clinicopathologic and structural considerations. Amyloid. 2003;10(Suppl. 1):48– 54.

[8] Yamada M. Cerebral amyloid angiopathy: an overview. Neuropathology. 2000;20:8–22.

[9] Benson MD. Amyloidosis. In: Scriver CR, Beaudet AL, Sly WS, et al, editors. The metabolic and molecular bases of inherited diseases. 8th ed. Vol. 4. New York (NY): McGraw-Hill; 2001. p. 5345–5378.

[10] Conceição I, Gonzalez-Duarte A, Obici L, et al. “Red-flag” symptom clusters in transthyretin familial amyloid polyneuropathy. J Peripher Nerv Syst. 2016;21:5–9.

[11] Shin SC, Robinson-Papp J. Amyloid neuropathies. Mt Sinai J Med. 2012;79:733–748.

[12] Kelly JW. Amyloid fibril formation and protein misassembly: a structural quest for insights into amyloid and prion diseases. Structure. 1997;5:595–600.

[13] Rowczenio DM, Noor I, Gillmore JD, et al. Online registry for mutations in hereditary amyloidosis including nomenclature recommendations. Hum Mutat. 2014;35:E2403–E2412.

[14] Hawkins PN, Ando Y, Dispenzeri A, et al. Evolving landscape in the management of transthyretin amyloidosis. Ann Med. 2015;47:625–638.

[15] Almeida MR, Damas AM, Lans MC, et al. Thyroxine binding to transthyretin Met 119. Comparative studies of different heterozygotic carriers and structural analysis. Endocrine. 1997;6:309–315.

[16] Hammarström P, Wiseman RL, Powers ET, et al. Prevention of transthyretin amyloid disease by changing protein misfolding energetics. Science. 2003;299:713–716.

[17] Sant’Anna R, Almeida MR, Varejao N, et al. Cavity filling mutations at the thyroxine-binding site dramatically increase transthyretin stability and prevent its aggregation. Sci Rep. 2017;7:44709.

[18] Sekijima Y, Dendle MT, Wiseman RL, et al. R104H may suppress transthyretin amyloidogenesis by thermodynamic stabilization, but not by the kinetic mechanism characterizing T119 interallelic trans-suppression. Amyloid. 2006;13:57–66.

[19] Terazaki H, Ando Y, Misumi S, et al. A novel compound heterozygote (FAP ATTR Arg104His/ATTR Val30Met) with high serum transthyretin (TTR) and retinol binding protein (RBP) levels. Biochem Biophys Res Commun. 1999;264:365–370.

[20] Coelho T, Chorao R, Sousa A, et al. Compound heterozygotes of transthyretin MET30 and transthyretin MET119 are protected from the devastating effects of familial amyloid polyneuropathy. Neuromuscular Disord. 1996;6(Suppl. 1):S20 Abstract 7.

[21] Coelho T, Carvalho M, Saraiva M, et al. A strikingly benign evolution of FAP in an individual found to be a compound heterozygote for two TTR mutations: TTR Met-20 and TTR Met-119. J Rheumatol. 1993;20:Abstract 179.

[22] Hammarström P, Schneider F, Kelly JW. Trans-suppression of misfolding in an amyloid disease. Science. 2001;293:2459–2462.

[23] Hammarström P, Jiang X, Hurshman AR, et al. Sequence-dependent denaturation energetics: a major determinant in amyloid disease diversity. Proc Natl Acad Sci USA. 2002;99(Suppl. 4):16427–16432.

[24] Bulawa CE, Connelly S, Devit M, et al. Tafamidis, a potent and selective transthyretin kinetic stabilizer that inhibits the amyloid cascade. Proc Natl Acad Sci USA. 2012;109:9629–9634.

[25] Hornstrup LS, Frikke-Schmidt R, Nordestgaard BG, et al. Genetic stabilization of transthyretin, cerebrovascular disease, and life expectancy. Arterioscler Thromb Vasc Biol. 2013;33:1441– 1447.

[26] Buniello A, MacArthur JAL, Cerezo M, et al. The NHGRI-EBI GWAS Catalog of published genome-wide association studies, targeted arrays and summary statistics 2019. Nucleic Acids Res. 2019;47:D1005–D1012.

[27] Allen N, Sudlow C, Downey D, et al. UK Biobank: current status and what it means for epidemiology. Health Policy Technol. 2012;1:123–126.

[28] Bycroft C, Freeman C, Petkova D, et al. The UK Biobank resource with deep phenotyping and genomic data. Nature. 2018;562:203–209.

[29] Bycroft C, Freeman C, Petkova D, et al. Genome-wide genetic data on ∼500,000 UK Biobank participants. BioRxiv. Available from: https://www.biorxiv.org/content/10.1101/166298v1 accessed September 5, 2019.

[30] O’Connell J, Sharp K, Shrine N, et al. Haplotype estimation for biobank-scale data sets. Nature Gen. 2016;48:817–820.

[31] Johnson JL, Abecasis GR. GAS power calculator: web-based power calculator for genetic association studies. BioRxiv. Available from: https://www.biorxiv.org/content/10.1101/164343v1 accessed September 5, 2019.

[32] Maurer MS, Schwartz JH, Gundapaneni B, et al. Tafamidis treatment for patients with transthyretin amyloid cardiomyopathy. N Engl J Med. 2018;379:1007–1016.

[33] Rosenblum H, Castano A, Alvarez J, et al. TTR (transthyretin) stabilizers are associated with improved survival in patients with TTR cardiac amyloidosis. Circ Heart Fail. 2018;11:e004769.

[34] Ando Y, Coelho T, Berk JL, et al. Guideline of transthyretin-related hereditary amyloidosis for clinicians. Orphanet J Rare Dis. 2013;8:31.

[35] Barroso FA, Judge DP, Ebede B, Li H, et al. Long-term safety and efficacy of tafamidis for the treatment of hereditary transthyretin amyloid polyneuropathy: results up to 6 years. Amyloid. 2017;24:194–204.

[36] Coelho T, Maia LF, Martins da Silva A, et al. Tafamidis for transthyretin familial amyloid polyneuropathy: a randomized, controlled trial. Neurology. 2012;79:785–792.

[37] Lozeron P, Theaudin M, Mincheva Z, et al. Effect on disability and safety of tafamidis in late onset of Met30 transthyretin familial amyloid polyneuropathy. Eur J Neurol. 2013;20:1539– 1545.

[38] Merlini G, Plante-Bordeneuve V, Judge DP, et al. Effects of tafamidis on transthyretin stabilization and clinical outcomes in patients with non-Val30Met transthyretin amyloidosis. J Cardiovasc Trans Res. 2013;6:1011–1120.

[39] Plante-Bordeneuve V, Gorram F, Salhi H, et al. Long-term treatment of transthyretin familial amyloid polyneuropathy with tafamidis: a clinical and neurophysiological study. J Neurol. 2017;264:268–276.

[40] Waddington Cruz M, Amass L, Keohane D, et al. Early intervention with tafamidis provides long-term (5.5-year) delay of neurologic progression in transthyretin hereditary amyloid polyneuropathy. Amyloid. 2016;23:178–183.

